# Genomic prediction in an outcrossing and autotetraploid fruit crop: lessons from blueberry breeding

**DOI:** 10.1101/2021.03.05.434007

**Authors:** Luís Felipe V. Ferrão, Rodrigo R. Amadeu, Juliana Benevenuto, Ivone de Bem Oliveira, Patricio R. Munoz

## Abstract

Blueberry (*Vaccinium corymbosum* and hybrids) is a specialty crop, with expanding production and consumption worldwide. The blueberry breeding program at the University of Florida (UF) has greatly contributed to the expansion of production areas by developing low-chilling cultivars better adapted to subtropical and Mediterranean climates of the globe. The breeding program has historically focused on phenotypic recurrent selection. As an autopolyploid, outcrossing, perennial, long juvenile phase crop, blueberry’s breeding cycles are costly and time-consuming, which results in low genetic gains per unit of time. Motivated by the application of molecular markers for a more accurate selection in early stages of breeding, we performed pioneering genomic prediction studies and optimization for implementation in the blueberry breeding program. We have also addressed some complexities of sequence-based genotyping and model parametrization for an autopolyploid crop, providing empirical contributions that can be extended to other polyploid species. We herein revisited some of our previous genomic prediction studies and described the current achievements in the crop. In this paper, our contribution for genomic prediction in an autotetraploid crop is three-fold: i) summarize previous results on the relevance of model parametrizations, such as diploid or polyploid methods, and inclusion of dominance effects; ii) assess the importance of sequence depth of coverage and genotype dosage calling steps; iii) demonstrate the real impact of genomic selection on leveraging breeding decisions by using an independent validation set. Altogether, we propose a strategy for the use of genomic selection in blueberry, with potential to be applied to other polyploid species of a similar background.

## 1 Introduction

Blueberry (Vaccinium corymbosum and hybrids) is recognized worldwide for its health benefits due to the high content and diversity of polyphenolic compounds (Kalt *et al.*, 2020). Such health-related attributes has resulted in an increased demand for blueberries, as it has become a crop with one of the highest production trends, with an increase of 142% of its production in the last 10 years (FAOSTAT, 2021). In this sense, the blueberry breeding program at the University of Florida (UF) has had a major contribution in the expansion of production areas. Starting in the 1950’s, the UF blueberry breeding program led pioneering hybridizations between high-quality US northern adapted species (*Vaccinium corymbosum*) and endemic US southern species (e.g., *Vaccinium darrowii*), selecting for low-chill requirements to break dormancy of flower buds (Sharpe and Sherman, 1971; Lyrene, 2000). The resulting breeding material and cultivars, known as southern highbush blueberries, established a new industry in Florida and in other warmer regions worldwide, allowing a year-round supply of fresh blueberries for the global market.

Historically, the UF program, like many others, used phenotypic recurrent selection with visual assessment of plants to select both new parents for crossing and clones to for commercial testing (Cellon *et al.*, 2018). Despite the success of the industry and the release of many cultivars in recent decades, the use of conventional methods results in low genetic gains per unit of time. Moreover, the autopolyploid nature of the crop, long juvenile phase, multi-year evaluations, large experimental areas, and the high sensibility to inbreeding depression makes phenotypic selection costly and time-consuming. Remarkably, it can take up to 12 years to release a new cultivar using conventional tools (Lyrene, 2005). As DNA sequencing costs continue to decrease, genomics-based markers present an opportunity to accelerate the breeding process by achieving more accurate selection during earlier breeding stages. Thus, the UF blueberry breeding program has been leading innovative genomics studies and procedures to fill two primary gaps in the blueberry breeding literature: understanding the genetic architecture of complex traits via genome-wide association studies (GWAS) and quantitative trait loci (QTL) mapping; and, at the practical level, performing phenotypic prediction based on molecular markers, a methodology popularly referred to as genomic selection.

GWAS and QTL mapping are both tools for providing biological elucidation of the genetic architecture, in which molecular markers spanning the entire genome are statistically tested for associations with phenotypes (Pritchard *et al.*, 2000). While QTL analyses are usually performed using structured populations, GWAS increases the mapping resolution by making use of populations with low levels of linkage disequilibrium considering a deep history of recombination events. In blueberry, we recently detected candidate genomic regions and markers associated with different fruit quality traits (FerrÃo *et al.*, 2018) and flavor-related volatiles (FerrÃo *et al.*, 2020) via GWAS investigations; and we built a high-density linkage map and detected QTL associated to berry firmness (Cappai *et al.*, 2020a). In counterpart, genomic selection aims to predict breeding values by using all genome-wide markers simultaneously (Meuwissen *et al.*, 2001). The underlying rationale is that, whenever the marker density is high enough, most QTL’s will be in linkage disequilibrium with some markers. Therefore, the estimated effect of all markers will lead to accurate predictions of the genetic merit for a complex trait. We have recently shown the potential of genomic selection in blueberry breeding under distinct modeling scenarios (de Bem Oliveira *et al.*, 2019, 2020; Amadeu *et al.*, 2020; Zingaretti *et al.*, 2020).

The autopolyploid nature of blueberry (2n=4X=48) imposes additional challenges for analysis and interpretation of genetic data. Autopolyploids possess genomes with multiple sets of homologous chromosomes, resulting in non-preferential pairing and potential polysomic inheritance during meiosis. Given the presence of higher allele dosage (i.e., the number of copies of each allele at a particular locus), a higher number of genotypic classes are possible (Gallais, 2003; Garcia *et al.*, 2013; Dufresne *et al.*, 2014). The inclusion of allelic dosage information on genomic selection models could imply in a more accurate estimation of breeding values by considering the additive effect of multiple copies of the same allele and the potential inheritance of dominance effects. However, accurate allele dosage calling on polyploids depends on higher depth of coverage which can increase genotyping costs when using sequence-based genotyping platforms (Gerard *et al.*, 2018; Caruana *et al.*, 2019). After performing foundational studies on the importance of polyploid models, inclusion of non-additive effects, and sequencing depth on allele dosage parameterizations, the UF blueberry breeding program is now on track to overcome the barrier of a simple promise to guide breeding decisions, to make genomic selection a reality.

Motivated by the potential to use genomic selection to reshape traditional blueberry breeding, we herein revisited some of our previous studies and described the current achievements in blueberry. Thus, our contributions in this paper are three-fold: (i) summarize previous results on the relevance of model parametrizations, such as diploid or polyploid methods, and inclusion of additive and non-additive gene actions for prediction; (ii) assess the importance of accurate dosage estimation for genomic prediction under low and high sequencing depth scenarios; (iii) demonstrate the realized impact of genomic selection over breeding cycles by using an independent validation set. Altogether, we anticipate elucidating challenges and directions for future studies in blueberry and relating our findings to other polyploid and fruit species with a similar breeding background.

## 2 Material and Methods

### 2.1 Populations and phenotypic data

The southern highbush blueberry populations used in this study were generated as part of the breeding program at the University of Florida. Two phenotypic datasets, referred as calibration set and testing set, were used with different purposes. The calibration set is comprised of a large breeding population already in use and is described in previous studies (FerrÃo *et al.*, 2018; de Bem Oliveira *et al.*, 2019). Briefly, it consists of 1,834 individuals originating from 117 biparental crosses using 146 distinct parents. All phenotypic evaluations were conducted on ripe fruits collected from the beginning of April to mid-May. Fruit firmness (g*mm −1 of compression force), size (mm), and weight (g) were evaluated over two seasons (2014 and 2015), while soluble solid (Brix) was evaluated only in 2015. Given the large representative population, all genomic prediction models reported in this study were calibrated using this dataset. The empirical best linear unbiased estimates (eBLUEs) were estimated for each genotype based on a linear model, where genotype and year were considered fixed effects, as described by Amadeu *et al.* (2020). Hereafter, the eBLUEs for each trait were considered as our response variable in the genomic prediction analyses.

The testing set was used for real validation in genomic prediction analyses. It comprises 280 advanced selections not originally included in the calibration set. These genotypes represent materials in the final evaluation stages in the breeding program. Hence, they were evaluated over several years in different locations throughout the state of Florida. As these phenotypes were collected from plants in different physiological phases and multiple environments, we adjusted the phenotypes using a linear model including separate fixed effects for year, location, and plant age. The eBLUEs of each genotype per trait were used as the phenotypic value in subsequent genomic prediction analyses. All phenotypic analyses were carried out using the ASREML-R software (Butler *et al.*, 2009).

### 2.2 Genotyping

The calibration set was genotyped using the “Capture-Seq” approach as described in Benevenuto *et al.* (2019). The genotyping of the testing set was also performed using “Capture-Seq” considering 10,000 biotinylated probes of 120-mer at RAPiD Genomics (Gainesville, FL, USA). Sequencing was carried out in the Illumina HiSeq2000 platform using 150 cycle paired-end runs. To ensure that the same group of single nucleotide polymorphisms (SNPs) will be called in both calibration and testing sets, we included the next-generation sequence data from both sets under the same SNP calling pipeline. Raw reads were cleaned and trimmed. The remaining reads were aligned using Mosaik v.2.2.3 (Lee *et al.*, 2014) against the largest scaffolds of each of the 12 homoeologous groups of *Vaccinium corymbosu*m cv. ‘Draper’ genome assembly (Colle *et al.*, 2019). Single nucleotide polymorphisms (SNPs) were called with FreeBayes v.1.3.2 using the 10,000 probe positions as targets (Garrison and Marth, 2012). Loci were filtered out applying the following criteria: minimum mapping quality of 10; only biallelic locus; maximum missing data of 50%; minor allele frequency of 1%; and minimum and maximum mean sequence depth of 3 and 750 across individuals, respectively. A total of 63,552 SNPs were kept after these filtering steps. Sequencing read counts per allele per individual were extracted from the variant call file using vcftools v.0.1.16 (Danecek *et al.*, 2011) and subsequently used to investigate some practical questions for the implementation of genomic prediction in polyploids.

Within the calibration set, we first investigated the importance of accurate genotype calling by comparing two strategies: (i) for the *ratio* method, each genotypic score was computed as the ratio between the alternative and total read depth, as described by SverrisdÓttir *et al.* (2017) and applied in de Bem Oliveira *et al.* (2019); (ii) for the *dosage method*, genotypic classes were assigned probabilistically using the updog v.2.1.0 R package considering the “norm model” and prior bias equals zero (Gerard *et al.*, 2018; Gerard and FerrÃo, 2020). Both genotyping methods (*ratio* and *dosage*) were compared under scenarios of high sequencing depth (random sampling for mean number of 60 reads – 60x) and low sequencing depth (random sampling for mean number of 6 reads – 6x). Specifically, we assumed the sequencing reads of each allele (alternative or reference) for a given marker come from a multinomial distribution, with probability equal the number of the reads divided by the total number of reads across all the alleles, markers, and individuals (N). Then, we sampled N/10 reads from this multinomial distribution. We performed this sampling 10 times, and each sampling result was used in a different cross validation fold. To avoid an eventual confounding between the number of markers and the predictive capacity over the four scenarios, we kept the same number of SNPs (63,552) across all scenarios. Therefore, in total, four scenarios were tested: *ratio* 60*x, ratio* 6*x, dosage* 60*x*, and *dosage* 6*x*.

For the real validation and implementation of genomic selection in the blueberry breeding program, we used the actual read counts to estimate the allele dosage in the calibration and testing sets according to the “norm model” in the updog v.2.1.0 R package (Gerard *et al.*, 2018; Gerard and FerrÃo, 2020). The posterior probability modes were used as our genotypic score. After estimating the posterior mean per genotype, we filtered out markers with a proportion of individuals genotyped incorrectly (*prop miss <* 10%), and markers with estimated bias higher than 0.13 and smaller than 7.38. Missing genotypes were imputed by the mean of each locus. A total of 48,829 SNPs were kept and used in genomic prediction for real validations.

### 2.3 Statistical Analyses

Single-trait linear mixed models were used to predict breeding values using the restricted maximum likelihood approach (REML) as following: *y* = *µ* + *Zu* + *e*; where **y** is a vector of pre-corrected phenotypic records for a particular trait; *µ* is the overall mean; **Z** is an incidence matrix linking observations in the vector y to their respective breeding value in the vector **u**. Normality where assumed for the additive and residual effects, where 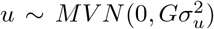 and the residual variance 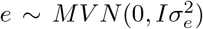. For the residual, **I** is an identity matrix; while and are the genetic and residual variance components. The matrix **G** denotes the genomic relationship matrix computed either using the *ratio* genotypic score or the tetraploid allele dosages with the different sequencing depths as described above. The matrices were estimated in the AGHmatrix v.2.0.0 R package (Amadeu *et al.*, 2016). For the *ratio* implementation, we used the “ratio” option in the software that compute the relationship as: 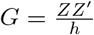, where **Z** is the mean-centered matrix of the molecular marker information (ratio values); and *h* is a scale factor, where 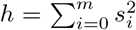 and 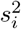 is the variance of the vector *z_i_* centered marker vector (for more details, see de Bem Oliveira *et al.* (2019)). For the *dosage* implementation, we used the additive relationship matrix based on VanRaden (2008) as described by de Bem Oliveira *et al.* (2019). All genomic prediction analyses were carried out using the rrBLUP package (Endelman, 2011).

Predictive performances were accessed for the *ratio* and *dosage* methods under high (60x) and low (6x) sequencing depth scenarios using only the calibration set in a 10-fold cross-validation scheme. To this end, the calibration set was randomly divided into 10 groups, where one group was used as a validation test, while the remaining nine groups were used as training. Models were trained in the validation test using the GBLUP approach. For each fold, predictive abilities were estimated using Pearson’s correlation between genomic estimated breeding values (GEBVs) and the corresponding eBLUEs. We also evaluated the correspondence between the top 20 group of individuals ranked using *dosage* 60*x* and the other scenarios. A *post hoc* Tukey test (p.value=.05) was used for intergroup comparisons between the top 20 ranked genotypes.

For the real genomic selection validation over the breeding cycles, we assessed the robustness of our predictive model over different stratification levels: (i) *General* predictions stand for models trained in the calibration set and predictions carried out in the testing data, in which the target phenotypic values were pre-corrected for year, location, and age fixed effects; (ii) For the *across-stages* predictions, a group of 114 individuals originally included in the calibration set but not in the testing set, was cloned in 2014 and planted in a commercial condition in a single location – prediction accuracy in this scenario can demonstrate the potential losses when models are trained at earlier stages and used at late stages of selection; (iii) In the *stratified* predictions, models trained in the calibration set were tested for predictions across four regions in Florida (North-FL, Central-FL, South-FL, and Citra-FL) – in contrast to the *general* predictions, in this scenario the target phenotypic values were pre-corrected only for the year effect per region. In all scenarios predictive performances were accessed via Person’s correlation between predicted and corrected phenotypic values, depending of the validation scenario.

A summary of all validation scenarios is illustrated in the Figure 1. For the stratified predictions, we complemented the predictive analysis by accessing the importance of genotype-by-environment interaction (GxE) via Analysis of Variance (ANOVA). To this end, we considered 16 genotypes (checks) that were phenotyped over the four regions. We fitted a linear model considering year, genotype, location, and the interaction between genotype and location (GxE) as fixed effects Analysis of Variance were performed in R (Team, 2013) using the native lm() function.

**Figure 1:**
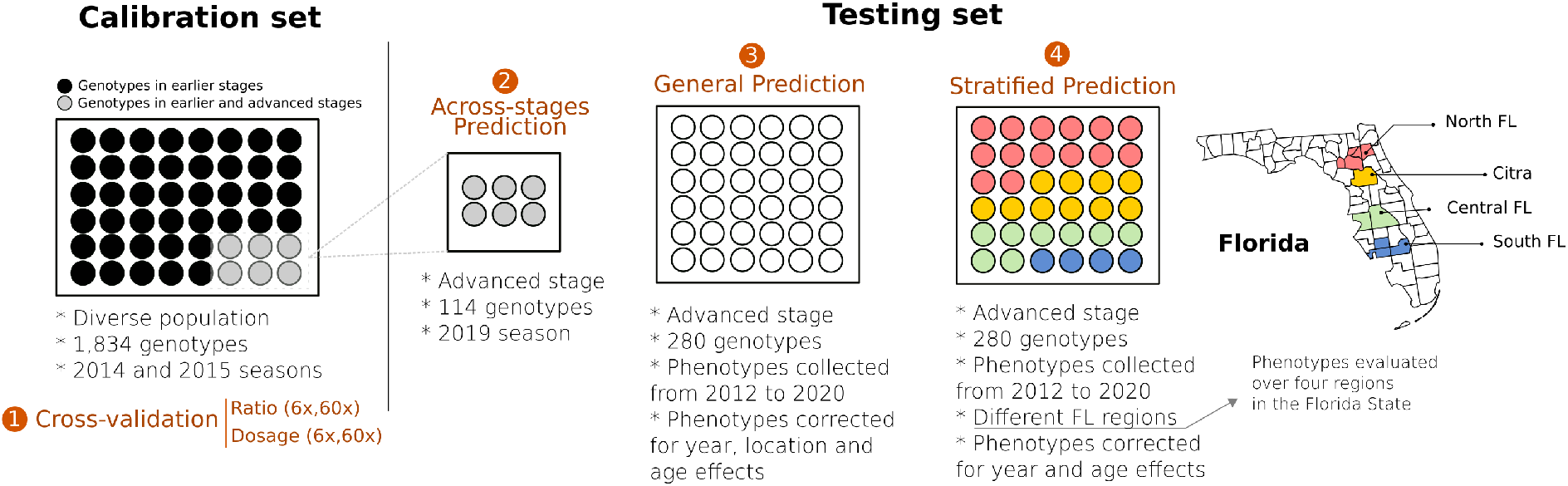
Schematic representation of four validation scenarios tested in blueberries. Calibration set represents a diverse group of genotypes were all genomic selection models where originally trained (de Bem Oliveira *et al.*, 2019). In the calibration set, using 10-fold cross validation, we tested the relevance of genotyping calling (*ratio* vs. *dosage*) under two different sequencing depth (6x and 60x) in scenario 1 of validation. Scenario 2 (*across-stages*) represents a group of 114 individuals originally presented in the calibration set that were clonally propagated, moved to advanced stage of the breeding program, and phenotyped under commercial field conditions. Scenario 3 (*general* prediction) represents a group of independent 280 genotypes, evaluated under commercial conditions, in which the phenotypic values of the target individuals were pre-adjusted for the year, location and age effects. Scenario 4 (*stratified* prediction) is an attempt to perform predictions over four regions of the State of Florida.

## 3 Results and Discussion

In the last two decades, genomic selection has become a reality for many animal and plant breeding programs. Despite the optimism and proved efficacy, its wide implementation is still hindered by investment costs and analytical skills required (Hickey *et al.*, 2017). With that in mind, the UF blueberry breeding program initiated genomic studies on a large scale in 2013. To begin, we worked closely with genotyping companies to design customized genotyping platforms; we phenotyped and genotyped a large and diverse blueberry breeding population; we increased our computational resources; and finally, we adapted our breeding framework to incorporate genomics. During this process, the implementation of genomic selection in polyploid and outcrossing species proved to be challenging, particularly regarding the intrinsic biological complexities and the availability of genomic and computational tools (Mackay *et al.*, 2019). In blueberry, for example, a high-quality genome assembly became available only in 2019 (Colle *et al.*, 2019). About half of the capture-seq genotyping probes that were originally developed based on a draft genome assembly were discarded afterwards based on the high-quality genome, without compromising genetic association and ge nomic prediction analyses (Benevenuto *et al.*, 2019). We also explored additional optimizations to reduce costs, regarding the number of individuals per family, number of markers, and sequencing depth (de Bem Oliveira *et al.*, 2020). Moreover, new genomics methods and tools have been developed in the last decade for the polyploid community, including allele dosage estimation, haplotype reconstruction, and the use of different relationship matrices (Bourke *et al.*, 2018). Here, we presented the lessons we have learned so far for implementing genomic selection in an autotetraploid and outcrossing species. We summarized previous results and also included novel findings relevant to the blueberry and polyploid community.

### 3.1 Filling the gaps: phenotypic and genotypic selection in the same breeding framework

Blueberry is an outcrossing and clonally propagated crop, for which the breeding process can be conventionally organized in two central steps: population improvement and product development (Lyrene, 2005). First, population improvement is done to manage the frequency of beneficial alleles over time by selecting and crossing outstanding materials, as conceptualized in recurrent selection designs. In parallel, product development consists of a series of trials in which potential candidates are evaluated over several years and locations, advancing across stages until the selection of the best clones becomes a registered variety. In Figure 2, we illustrated these two key steps and how they are integrated in a four-stage selection design (from Stage I to IV) in the UF blueberry breeding program.

**Figure 2:**
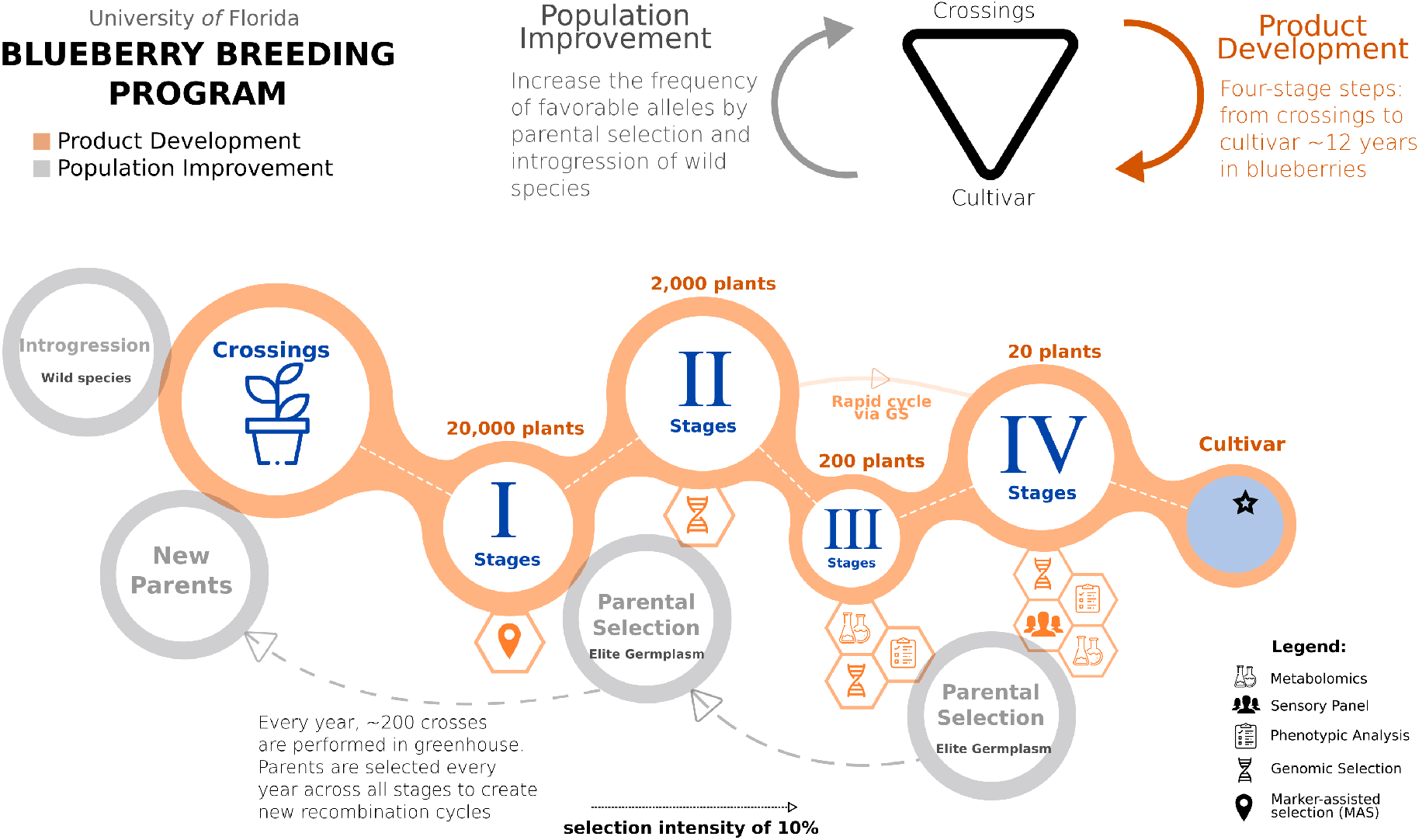
A schematic representation of the UF blueberry breeding program, integrating phenotypic and genomic prediction. The breeding process is conventionally organized in two integrated steps: population improvement (orange) and product development (gray). A breeding cycle starts with crosses between outstanding parental genotypes. After that, several stages (I to IV) are required to evaluate the genotype performance. At Stage I, we will use marker-assisted selection targeting traits with simple genetic architecture. Genomic selection will be implemented in Stages II, when GEBVs are computed. In advanced selections (Stages III and IV), high-quality phenotyping will be performed to leverage the calibration of genomic prediction models. At these stages, the use of metabolomics and sensory panel analyses will also play an important role for flavor-assisted selections. At the end, elite materials are registered as clonally propagated cultivars. Rapid cycles could be achieved by selecting plants directly from the Stages II to Stages IV, as originally proposed by de Bem Oliveira *et al.* (2019). For population improvement, over the four-stage design, elite germplasms and wild species are systematically selected to constitute new breeding cycles.

Annually, the blueberry breeding program performs more than 200 crosses, including parents selected among cultivars, elite material, and wild germplasm (Lyrene, 2005). From these crosses, about 20,000 seedlings are planted in non-replicated high-density nurseries, establishing the so-called Stage I. After one year, plants in the Stage I are visually selected based on fruit size, color, scar, and using the breeder’s ‘bite test’ for flavor quality attributes. Approximately 10% of the original number of seedlings are kept after this first selection. To not exhaust genetic diversity, a minimal number of individuals per family are kept. Given blueberry’s long juvenile period, the availability of few berries, and the high competition in a high-density planting, it is difficult to phenotype for all traits and assess the full potential of the individuals at this stage. Additionally, the large number of individuals prevents the use of genomic prediction at this stage, given the costs of genotyping. Therefore, at Stage I, we envision that the use of marker-assisted selection (MAS) for traits with simple genetic architecture is a more feasible approach, and it is a current research line of the breeding program. In this regard, example of MAS implementation in early selection stages are reported in strawberry (Gezan *et al.*, 2017; Osorio *et al.*, 2020).

After the first selection, approximately 2,000 genotypes pass to the second stage (Stage II). All plants stay in the same field plot, while unselected ones are removed. Further visual phenotypic evaluations are performed for the next 3 years. It is at this stage that we are implementing genomic prediction to increase genetic gains by improving phenotyping accuracy and selecting parents at early stages. Therefore, at Stage II, all plants will be genotyped, and the GEBVs will be predicted for five fruit quality traits (soluble solids, titratable acidity, weight, size, and firmness), yield, and consumers panel liking scores. Using a selection index according to trait importance, we will perform genomic selection to complement standard phenotypic descriptors and rank all genotypes. As routinely done, 10% of the 2,000 plants will be moved to the next stage (Stage III), where plants are clonally propagated and evaluated in a 15-plant clonal plot in a commercial field.

At Stage III, around 200 plants are more accurately phenotyped for more traits, using more fruits, clonal repetitions, and multiple years of evaluations in commercial conditions. Technically, all information collected at this stage will be used to feed the genomic prediction models. In recent years, the UF blueberry breeding program has included new traits for routine phenotyping to meet the current demand from different marketable demands. For example, the use of volatiles for flavor-assisted selection has shown the ability to predict sensory perceptions, and thus, metabolomics have been incorporated in the breeding pipeline to assess flavor ratings for a large number of genotypes (Gilbert *et al.*, 2014; Colantonio *et al.*, 2020; FerrÃo *et al.*, 2020). Similarly, we want to incorporate nutraceutical compounds to leverage blueberry health benefits and shelf-life related traits so berries can withstand long periods of storage or shipment.

In the last stage (Stage IV), around 15-20 plants selected from Stage IIIs with consistent and outstanding performances are propagated and planted at commercial trial sites across the state of Florida. The different locations comprise two production systems according to the accumulation of chilling hours: evergreen and deciduous (Fang *et al.*, 2020). To ensure accurate selection, phenotypic data is collected weekly and will also be used to feed our genomic prediction models. Fruits from selected genotypes are also submitted to sensory panels, where flavor preferences are scored by blueberry consumers. Elite selections from this final stage are ultimately named, patented, and released as clonally propagated cultivars.

Altogether, the conventional breeding pipeline takes up to 12 years to evaluate the genotype merit of an individual to be released as a cultivar. With the implementation of genomic prediction as the scope of the breeding program, the selection criteria will be more accurate than visual phenotypic selection at the Stage II. Moreover, it will shorten the time to select genotypes to advance to Stage III and to become a parent in the next breeding cycle. In a typical recurrent selection breeding scheme, parental selection is a crucial step (Lyrene, 2005). We have optimized this selection not only by ranking the GEBVs over the breeding cycles, but also by seeking crosses that minimize inbreeding. Among the different tools available for mate allocation, we have recently implemented the algorithm described in the AlphaMate software (Gorjanc and Hickey, 2018).

### 3.2 “Simplicity is the ultimate sophistication^1^”: on the relevance of additive GBLUP models

When confronting the problem of modeling the relationship between molecular markers and variation in the observed traits, an important question to keep in mind is what statistical method could better describe this relationship (FerrÃo *et al.*, 2019). In recent years, we have investigated statistical and biological aspects underlying the implementation of genomic prediction in autopolyploid species, including (i) the importance of accounting for allele dosages in whole-genome statistical models (de Bem Oliveira *et al.*, 2019); (ii) the relevance of multiple gene actions, including additive and non-additive genetic sources (Amadeu *et al.*, 2020; Zingaretti *et al.*, 2020); and finally, (iii) the impact of sequencing depth of coverage, when sequence-based genotyping approaches are used (de Bem Oliveira *et al.*, 2020).

Among the factors that differentiate diploid and polyploid analyses, resolving the allelic dosage of individual loci is one the most important. While in diploid organisms, only three genotypic classes are possible for biallelic markers; autotetraploids, like blueberry, can have up to five genotypic classes. Therefore, in theory, it is expected that statistical models accounting for the dosage effect could be more informative and provide a more realistic representation of the genetic complexity of a quantitative trait (Garcia *et al.*, 2013). We first tested this hypothesis by contrasting polyploid and diploid parametrizations in GWAS studies (FerrÃo *et al.*, 2018); whereby, in fact, a larger number of associations were observed under polyploid models. In a subsequent study, we investigated a similar assumption for genomic prediction (de Bem Oliveira *et al.*, 2019) and tested GBLUP models using relationship matrices built in a tetraploid (Slater *et al.*, 2016) and diploid (VanRaden, 2008) fashion. Interestingly, both parametrizations resulted in similar performances for all traits tested. Similar predictive ability for diploid and polyploid parametrizations were also reported in other autotetraploid species (Lara *et al.*, 2019; Matias *et al.*, 2019), which ultimately reinforced the robustness of the predictive capacity of GBLUP regardless of the ploidy parametrization used.

Besides the potential additive impact of allele dosages, dominance effects can also be heritable in polyploids and could improve the prediction of genetic values. Therefore, it is also reasonable to speculate that a greater number of alleles per locus may increase the range of genetic models to describe one-locus genotypic value by accounting for multiple dominance levels (Gallais, 2003). This is exemplified by the different models addressing the dominance effect proposed in the polyploid literature, including the use of digenic interactions (Endelman *et al.*, 2018), the use of a general effect by assuming that each genotype has its own effect (Rosyara *et al.*, 2016; Slater *et al.*, 2016), and the use of heterozygous parametrization (Enciso- Rodriguez *et al.*, 2018). In blueberries, we tested the importance of such different gene action in predictive studies. Although we have observed an improvement in the statistical goodness-of-fit when dominance effects are counted, this increment is not directly translated into predictive ability (Amadeu *et al.*, 2020). Hence, the additive model resulted in performance similar to models accounting for dominance effects, as it has been described for diploid species (MuÑoz *et al.*, 2014).

Given the genetic complexity of polyploids and the potentially higher levels of intra- and inter-locus interactions, we also hypothesized that predictions could be improved by using deep learning techniques (Zingaretti *et al.*, 2020). Through deep learning, we could take advantage of non-linearity assumptions to model the whole genetic merit of an individual. To test this, we used allo-octoploid strawberry and auto-tetraploid blueberry as our biological models and compared linear models and deep learning techniques for prediction. In both species, we did not observe improvements of deep learning over traditional linear models for traits with presumably different genetic architectures. The only exception was observed in a simulated data set, in which deep learning performed better for traits with large epistatic effects and low narrow-sense heritability. This again, reinforced the high predictive capacity of mixed models as prediction machinery.

Our last contribution for the practical implementation of genomic prediction in polyploids is regarding the relevance of sequencing depth of coverage for genotyping methods based on next-generation sequencing. Sequencing depth refers to the number of reads sequenced at a given site in the genome. Low coverage datasets increase the chances of not sampling all homologous chromosomes at a given site for a given individual during sequencing. It could result in high rates of missing data, miscalled genotypes, and uncertainty of allele copy number in heterozygous genotypes (Clark *et al.*, 2019). To circumvent this issue, some studies in polyploid crops have recommended increasing the sequencing depth, which implies higher costs of genotyping. For example, Bastien *et al.* (2018) and Uitdewilligen *et al.* (2013) suggested sequencing depths of 50X–80X for an accurate assessment of allele dosage in autotetraploid potatoes. In a recent study, we demonstrated that such numbers are quite conservative for genomic prediction. By combining a simple genetic parametrization (*ratio*) and low-to-mid sequencing depth (6x-12x), we achieved similar predictive accuracies as higher-depths for blueberry traits with different genetic architectures (de Bem Oliveira *et al.*, 2020). Similar results are also reported by Zheng *et al.* (2020). In practical terms, reducing the amount of sequencing data will also reduce the costs associated with implementing genomic selection or potentially genotyping more individuals under a fixed budget.

Despite the considerable advancements previously explored, the relevance of using more sophisticated algorithms for genotype calling and its impact on genomic prediction remains unexplored. Recently, several new methods have been developed to assign accurate allelic dosage of individual loci in polyploids (Garcia *et al.*, 2013; Gerard *et al.*, 2018; Pereira *et al.*, 2018; Clark *et al.*, 2019). In this paper, we compared predictive abilities using different genomic parametrization and confirmed that low-to-mid sequencing depth and *ratio* parametrization can be used for ranking GEBVs – with similar predictive performance (Table 1) and genotypic ranking (Table 2). Nonetheless, despite the attractive simplicity of using the *ratio* and low-sequencing depth, such results are only valid for prediction studies (de Bem Oliveira *et al.*, 2019, 2020). Importantly, there is no empirical evidence that setting the parameters to these levels could work for inferential studies such as GWAS, population genomics, linkage and QTL mapping. In this sense, an important counterpoint was recently reported in hexaploid sweet potato, for which higher sequencing depths and accurate dosage calling improved the ultra-dense linkage map and posterior QTL analysis (Gemenet *et al.*, 2020; Mollinari *et al.*, 2020). For GWAS, we observed large rates of false positive associations when analyses were performed using low sequencing depth associated to the *ratio* parametrization (results not shown). Herein, we systematically observed large biases when relationship matrices were constructed using the *ratio* 6*x* approach (Figure S1 and Table S2).

**Table 1:** Predictive ability between two different genotype calling approaches (*dosage* and *ratio*) under two sequencing depth scenarios (6x and 60x) for five fruit quality traits in blueberry using 10-fold cross-validation. Results are means and, between brackets, the range observed.

| Method | Depth | Firmness | Size | Weight | Brix |
| --- | --- | --- | --- | --- | --- |
| *Dosage* | 6x | 0.44 [0.40-0.52] | 0.38 [0.29-0.49] | 0.47 [0.40-0.57] | 0.28 [0.18-0.47] |
| *Dosage* | 60x | 0.46 [0.41-0.55] | 0.40 [0.29-0.53] | 0.49 [0.40-0.56] | 0.29 [0.17-0.46] |
| *Ratio* | 6x | 0.45 [0.38-0.55] | 0.40 [0.31-0.48] | 0.48 [0.42-0.56] | 0.28 [0.18-0.43] |
| *Ratio* | 60x | 0.46 [0.40-0.54] | 0.39 [0.29-0.51] | 0.49 [0.41-0.56] | 0.29 [0.17-0.47] |

**Table 2:** Number of clones matching the top 20 ranking of clones using the *dosage* 60*x* method as benchmark, under 10-folds cross-validation. A *post hoc* Tukey test (alpha=.05) was used for intergroup comparisons over the scenarios. Cells with the same letter represent non-statistically different groups for the given trait (column).

| Method | Depth | Firmness | Size | Weight | Brix |
| --- | --- | --- | --- | --- | --- |
| *Dosage* | 6x | 16.5<sup>b</sup> | 16.9<sup>b</sup> | 16.3<sup>b</sup> | 16.4b |
| *Ratio* | 6x | 16.2<sup>b</sup> | 15.6<sup>c</sup> | 16.2<sup>b</sup> | 15.3b |
| *Ratio* | 60x | 18.8<sup>a</sup> | 18.3<sup>a</sup> | 18.6<sup>a</sup> | 18.7a |

In summary, our results are suggesting that the use of traditional GBLUP is robust enough for genomic prediction, even under the simplistic assumptions.This fact that has long been discussed in the specialized literature, and has raised questions on the contribution of linkage disequilibrium between QTLs and markers versus the relationship information to genomic selection (Habier *et al.*, 2013).

### 3.3 How does genomic prediction work in a real validation population?

While we have investigated several statistical and computational aspects related to genomic selection in blueberry, it is still unknown how accurate the predictions will be across breeding cycles, with plants in different phenological stages and locations. This scenario came to be called “true validation” and involves the use of independent populations. We investigate it by dividing our prediction analyses as following: models calibrated in 2014 and 2015 using plants in Stage II were used for phenotypic predictions of Stages III and IV individuals. Both data sets share genetic similarity (Figure 3 a).

**Figure 3:**
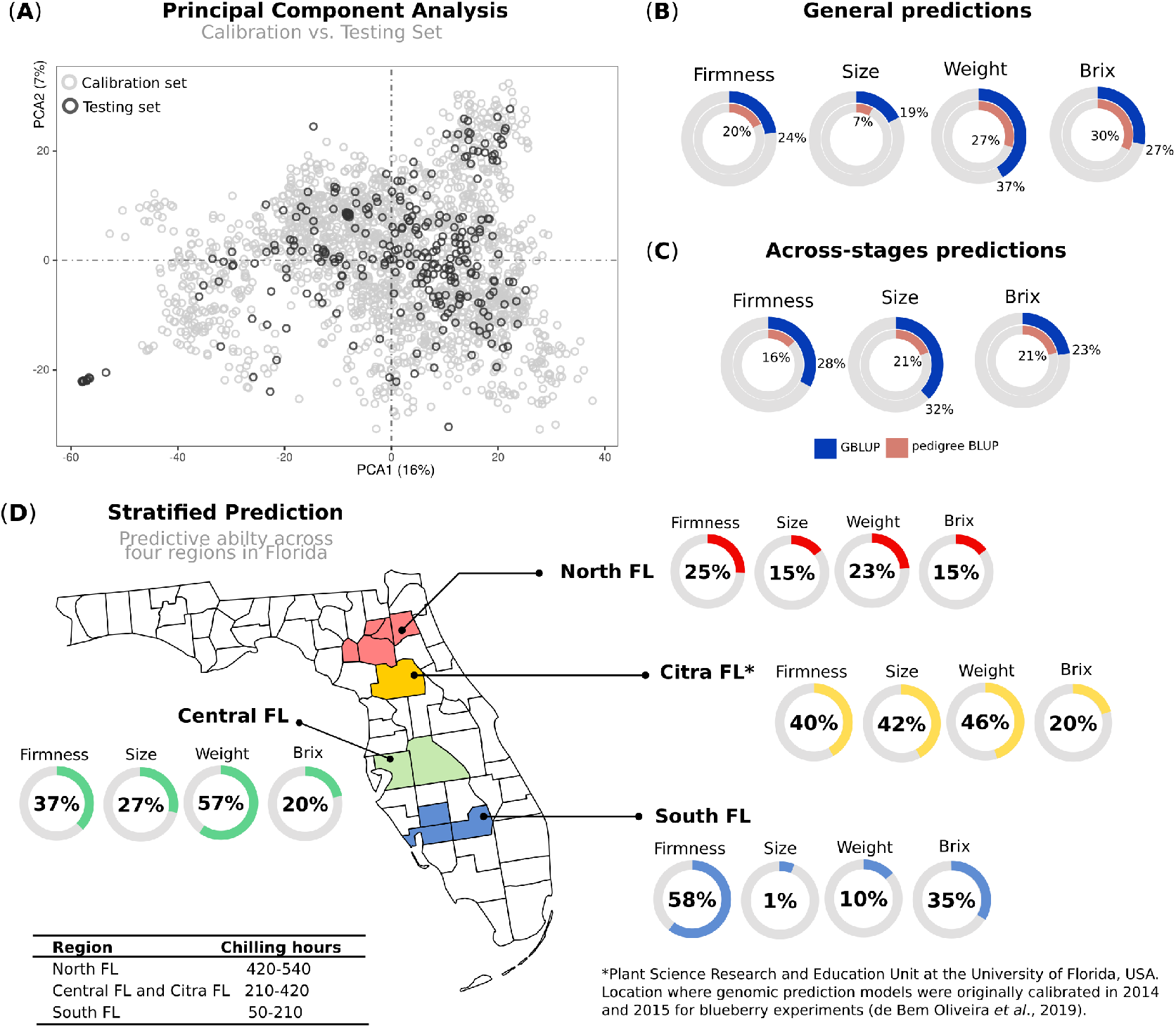
Phenotypic prediction. (A) Principal component analysis (PCA) of two blueberry populations: calibration set represents the trained set, where genomic prediction models were originally calibrated, and testing set comprising additional 280 individuals used for testing. (B) *General* prediction: predictive ability of the testing set, after training the models in the calibration set. (C) *Across-stages* prediction: predictive ability measured in a group of 114 individuals that was also included in the calibration set, but phenotypes were collected in advanced selection stages. (D) *Stratified* predictions: after training the models in the calibration set, individuals in the testing set were predicted using phenotypes collected over four macro-regions in the Florida State, which are under different chilling hour accumulation.

For true validations, we tested different scenarios in which genomic selection could be applied (Figure 1). First, we predicted the overall performance using genomic and pedigree information and confirmed the importance of genomic information (*general* predictions, in Figure 3b). When compared to predictions using within-sample cross-validation schemes, as reported in de Bem Oliveira *et al.* (2019) and Amadeu *et al.* (2020), we observed a reduction on the predictive results (Table S1). This decline in predictive performance in real validation is expected, due to differences in the allele frequencies over populations, variation in linkage disequilibrium patterns, and genotype-by-environment interactions (Habier *et al.*, 2013).

Another predictive scenario focused on validations across breeding cycles. To this end, we used the calibration test – originally evaluated in Stages II – to predict a subgroup of individuals that were cloned and planted in an advanced stage (Stages III). On average, larger values were observed compared to the general scenario, but it was still, as expected, lower than within-sample cross-validation schemes (*across- stages* predictions, Figure 3c). These results highlight (i) the importance of collecting better phenotypic data and (ii) the influence of the plant management. Remarkably, most of the phenotypic traits measured in the calibration set were collected from five berries per genotype; while on Stage III, we used 25 berries per genotype. Furthermore, genotypes in Stage II are planted in high-density nurseries with phenotypes collected in plants that are still in their juvenile phase, while Stages III are grown under commercial conditions.

In the third scenario, a more challenging exercise was to measure how predictive capacity varies across regions in the State of Florida (*stratified* predictions, Figure 3d). Higher predictability was observed for Citra and Central-FL, the closest regions where the models were originally trained. In counterpart, plants evaluated in the South-FL showed, on average, lower predictability performances. Despite the small number of genotypes included in this analysis, these results provide insights into the importance of genotype-by-environment (GxE) interaction for genomic selection in blueberry. We further explored this hypothesis by using a group of 16 common genotypes (checks) evaluated over the four regions. The results confirmed the significance of the GxE effect for most of the traits (Table 3), with the plants evaluated in South-FL showing the most contrasting values. It is noteworthy that blueberry locations in South-FL are grown under an evergreen production system, under less chilling hours, and are focused on preventing defoliation during the winter months (Fang *et al.*, 2020). On the other hand, the location in Citra, Central-FL, and North-FL regions are grown under the deciduous production system, where leaves are dropped during the winter. Such differences in the production systems could be driving the largest disparity observed at South-FL, when compared to the other regions.

**Table 3:** Mean and standard deviation (in parenthesis) of four fruit quality traits evaluated in advanced stages of the blueberry breeding program at four regions of Florida. Values were computed using 16 common genotypes (checks).

| Location | Firmness | Size | Weight | Brix |
| --- | --- | --- | --- | --- |
| North FL | 248 (32.8) | 18.0 (1.79) | 2.57 (0.641) | 11.3 (1.31) |
| Citra FL | 245 (42.0) | 17.0 (2.22) | 2.34 (0.691) | 10.9 (1.33) |
| Central FL | 244 (29.3) | 17.7 (1.46) | 2.29 (0.491) | 11.8 (1.27) |
| South FL | 251 (33.4) | 17.4 (1.34) | 2.21 (0.549) | 12.0 (1.91) |
| GxE (p.value)* | 0.007 | 0.0002 | 0.005 | 0.47 |

\* P.values associated to genotype-by-environment interaction (GxE) were computed using a linear model and Analysis of Variance (ANOVA), where season, genotype, location, and the interaction between genotype and location (GxE) were fitted as fixed effects.

The results from real validations allow us to draw some practical conclusions. First, even with low-to-moderate predictive accuracies, genomic selection is still encouraging. For example, soluble solids and firmness are both traits treasured by consumers, for which routine phenotyping is expensive and time-consuming for large populations, like Stage IIs. Ranking plants based on their GEBVs proved to be a better alternative than any other criteria historically used over the course of UF blueberry breeding program (pedigree or visual selection). More accurate phenotypic data to annually re-calibrate the model has also the potential to improve predictability. Finally, we also reinforce the importance to recalibrate our models considering the environment targets. With the recent advent of high-throughput phenotyping, we envision that more data across different production systems in Florida – and around the globe– could be used for better calibration and ultimately, more accurate predictions. Examples of image analyses for high-throughput phenotyping have been reported in other fruit trees (Diaz-Garcia *et al.*, 2016; Di Gennaro *et al.*, 2019; Koirala *et al.*, 2019; Feldmann *et al.*, 2020), including blueberries (Jiang *et al.*, 2019).

### 3.4 Unifying biological discoveries and predictions

The use of genomics information also provides new opportunities to integrate biotechnology and quantitative genetics into modern breeding programs, creating platforms for both delivery of new products and biological discovery (Hickey *et al.*, 2017). In blueberry, biological discoveries have been addressed via QTL mapping (Cappai *et al.*, 2020a) and GWAS studies (FerrÃo *et al.*, 2019, 2020) for multiple fruit quality traits. Unifying such new discoveries with prediction is challenging, but it has been addressed under three different avenues: (i) use of GWAS discovered QTLs as fixed effects on GS models; (ii) incorporating markers (or QTL) in MAS designs; and (iii) using genome-editing technology to speed up breeding.

In a strategy called “GS *de nov* o GWAS”, we explored the importance and applicability of GWAS findings for prediction by using the significant GWAS hits as fixed effects in GS models, considering independent datasets. For oligogenic traits, like some flavor-related volatiles, we achieved an increase of more than 20% in the predictive ability, when compared with traditional GS methods (FerrÃo *et al.*, 2018). Using a similar strategy, gains in predictive performance have been also reported in other crops, such as maize (Bernardo, 2014) (Rice and Lipka, 2019), wheat (Sehgal *et al.*, 2020), and rice (Spindel *et al.*, 2016). Alternatively, we have investigated further modelling strategies to accommodate biological information into the predictive models. For example, the use of Bayesian strategies that could accommodate SNPs with larger effect by using different prior distributions (Gianola, 2013; Zhou *et al.*, 2013; Erbe *et al.*, 2012); and GBLUP models that could weight variants previously selected either via association analysis or using bioinformatic pipelines (Su *et al.*, 2014; Zhang *et al.*, 2016; Liu *et al.*, 2020; Ren *et al.*, 2021).

Another potential strategy is to use target markers associated with important traits for MAS during Stage I of the blueberry breeding program. Such a strategy could be used for early selection of plants still in the seedling stage. Acknowledged by their simple genetic architecture, we showed that few markers could yield reasonable predictive accuracies of volatile emission and, thus, leverage flavor selection (FerrÃo *et al.*, 2020). We envision that MAS can also be implemented for other oligogenic traits. In this regard, we have been conducting other GWAS and QTL mapping studies for disease resistance, such as anthracnose (*Colletotrichum gloeosporioide*s) and bacterial wilt (*Ralstonia solanacearum*). A similar strategy has been implemented in strawberry (Gezan *et al.*, 2017; Osorio *et al.*, 2020) and other fruits (Iezzoni *et al.*, 2020). However, for MAS to be applicable for thousands of plants, fast DNA extraction and SNP genotyping assays should be optimized.

Gene editing is another attractive technology with potential to have significant effects in the breeding program. Aside from the use of CRISPR-Cas9 for validating candidate genes identified via GWAS or QTL studies, some simulations have recently shown that genome editing can double the rate of genomic gain when coupled with genomic prediction, compared to genomic selection conducted in isolation (Noman *et al.*, 2016; Hickey *et al.*, 2017). To our knowledge, there is only one study of CRISPR-Cas9 targeted mutagenesis in blueberry (Omori *et al.*, 2021). At the UF blueberry breeding program, we have advanced our understanding on the best tissue culture practices and most effective transformation markers (Cappai *et al.*, 2020b), laying the ground for CRISPR/Cas9 genome editing implementation in our breeding program. Using this technique, we can also take advantage of the knowledge accumulated from model crops to introduce novel allelic diversity in orthologues and accelerate the domestication process.

## 4 Conclusions ans Future Directions

The implementation of genomic selection has already changed the UF blueberry breeding program routine, by reorganizing the way we collect genotype and phenotype information and analyze data to rank the material to advance stages and to breed in the next cycles. Our previous studies on genomic selection were fundamental to define the most cost- and time-effective methods for model parameterization and genotyping. The main lessons learned can be conveniently divided in different areas. Statistically, despite the numerous algorithms for prediction – many of them more elegant at the biological and computational level – it was the use of additive effects under a linear mixed model framework (GBLUP) that showed the best balance between efficiency and accuracy. Considering the particularities of autopolyploid genetic data, we showed that for genomic selection, low depth of sequencing (6x-12x) and simplifying the allele dosage information (i.e., diploidization and *ratio*) resulted in similar prediction accuracies as those obtained using more refined scenarios. At the practical level, genomic prediction was incorporated in a recurrent selection breeding scheme, whereby variety deployment and populational improvement run in parallel. So far, GEBVs have been primarily used for parental selection to increase genetic gains, while keeping the genetic diversity.

Finally, we highlight some challenges and potential opportunities for further studies in blueberries. First, re-calibrating the model with more accurate phenotypic data can yield better predictive ability. In this sense, phenomics is also a cutting-edge area of research that could leverage the number of samples collected during a season and improve the quality of phenotypic data. For example, yield is a complex and time-consuming trait to be phenotyped over the season. We envision that image-based phenotyping may aid on the task of evaluating yield and other traits associated to plant architecture and diseases. Statistically, testing new algorithms for mate allocation, and using haplotypes for prediction and imputation methods are some potential areas that could further improve genomic predictions.

## Acknowledgements

The authors thank Catherine Cellon and James Olmstead for the calibration set data collection; Werner Collante, Lauren Scott, and Douglas Phillips for the testing set data collection; and Mia Acker for reviewing the manuscript. This work was supported by the UF royalty fund generated by the licensing of blueberry cultivars.

## Author contributions

PRM and LFVF conceived and supervised the study. JB coordinated the collection and genotyping of the samples. IBO coordinated the collection the data for real validation. LFVF and RRA analyzed and interpreted the phenotypic and genomic selection results. LFVF wrote the paper and included the revision from all authors. All authors read and approved the final version of the manuscript for publication.

## Abbreviations

UF: (University of Florida)
GEBV: (genomic Estimated breeding value)
GWAS: (genome-wide association studies (GWAS)
QTL: (quantitative trait loci)
EBLUE: (empirical best linear unbiased estimates)
SNP: (single nucleotide polymorphisms)
(GxE): genotype by environment interaction
MAS: (marker-assisted selection)

## Footnotes

1 Quote by Leonardo da Vinci

