## Supplemental Table 1 for "Genomic prediction in an outcrossing and autotetraploid fruit crop: lessons from blueberry breeding"

### *Supplementary Material*

**Table S1:** Predictive ability of calibration set previously reported in blueberry for fruit quality, production, and volatile organic compounds (VOCs) using tetraploid additive GBLUP.

| Trait | Category | Seasons | Population Size | Predictive ability | Reference |
|---|---|---|---|---|---|
| Soluble Solid | Fruit quality | 2014 | 1,847 | 0.28 | (de Bem Oliveira et al., 2019) |
| Size | Fruit quality | 2014,2015 | 1,847 | 0.40 | (de Bem Oliveira et al., 2019) |
| Firmness | Fruit quality | 2014,2015 | 1,847 | 0.46 | (de Bem Oliveira et al., 2019) |
| pH | Fruit quality | 2014 | 1,847 | 0.26 | (de Bem Oliveira et al., 2019) |
| Fruit Scar | Fruit quality | 2014,2015 | 1,847 | 0.47 | (de Bem Oliveira et al., 2019) |
| Fruit Weight | Fruit quality | 2014,2015 | 1,847 | 0.44 | (de Bem Oliveira et al., 2019) |
| Yield | Production | 2014,2015 | 1,847 | 0.35 | (de Bem Oliveira et al., 2019) |
| Flower Buds | Production | 2015 | 1,847 | 0.18 | (de Bem Oliveira et al., 2019) |
| (E)-2-Hexenal | VOCs | 2015 | 886 | 0.47 | (Ferrão et al., 2020) |
| 1-Hexanol | VOCs | 2015 | 886 | 0.46 | (Ferrão et al., 2020) |
| 2-Heptanone | VOCs | 2015 | 886 | 0.58 | (Ferrão et al., 2020) |
| 2-Nonanone | VOCs | 2015 | 886 | 0.67 | (Ferrão et al., 2020) |
| 2-Undecanone | VOCs | 2015 | 886 | 0.65 | (Ferrão et al., 2020) |
| D-limonene | VOCs | 2015 | 886 | 0.56 | (Ferrão et al., 2020) |
| Decanal | VOCs | 2015 | 886 | 0.56 | (Ferrão et al., 2020) |
| Eucalyptol | VOCs | 2015 | 886 | 0.49 | (Ferrão et al., 2020) |
| Geranyl acetone | VOCs | 2015 | 886 | 0.53 | (Ferrão et al., 2020) |
| Hexanal | VOCs | 2015 | 886 | 0.41 | (Ferrão et al., 2020) |
| Linalool | VOCs | 2015 | 886 | 0.53 | (Ferrão et al., 2020) |

**Table S2:** Person's correlation between off-diagonal values of genomic relationship matrices computed using dosage and ratio parameterizations under two sequencing depth scenarios (6x and 60x)

| | **dosage_60x** | **ratio_60x** | **dosage_6x** | **ratio_6x** |
|---|---|---|---|---|
| **dosage_60x** | 1.00 | 0.99 | 0.97 | 0.79 |
| **ratio_60x** | | 1.00 | 0.96 | 0.79 |
| **dosage_6x** | | | 1.00 | 0.83 |
| **ratio_6x** | | | | 1.00 |

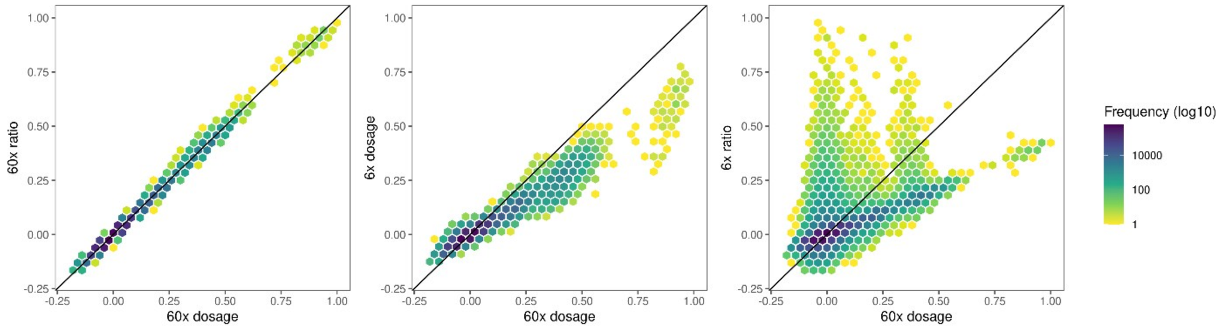


**Figure S1:** Scatterplot of off-diagonal values for the genomic relationship matrices computed using dosage and ratio parametrizations under two sequencing depth scenarios (6x and 60x). Diagonal line represent the absolute similarity between scenarios.
